# Benchmarking analysis of deleterious SNP prediction tools on CYP2D6 enzyme

**DOI:** 10.1101/760298

**Authors:** Karla Cristina do Vale Ferreira, Leonardo Ferreira Fialho, Octávio Luiz Franco, Sérgio Amorim de Alencar, William Farias Porto

## Abstract

The cytochrome P450 family is composed of hemeproteins involved in the metabolic transformation of endogenous and exogenous substances. The CYP2D6 enzyme is responsible for the metabolism of approximately 25% of clinically used drugs and is mainly expressed in the liver. The *CYP2D6* gene is known to have a large number of Single Nucleotide Polymorphisms (SNPs) and the majority of them do not present clinical consequences. Nevertheless, these variations could modify the CYP2D6 enzyme’s function, resulting in poor metabolizing or ultra-extensive metabolizing phenotypes, when metabolism is slower or accelerated, respectively. Currently, there are several computational tools for predicting functional changes caused by genetic variations. Here, using 20 web servers, we evaluated the impact of 21 missense SNPs (6 neutral and 15 deleterious) previously validated by the literature. Only seven predictors presented sensitivity higher than 70%, while four showed specificity higher than 70% and only one reached the Matthews correlation coefficient of 0.39. Combinations of tools with greater sensitivity and specificity were made to improve the Matthews correlation coefficient, which increased the coefficient of five tools (Provean, FatHMM, SDM, PoPMuSiC and HotMuSiC). The results suggest that the most appropriate tool for CYP2D6 SNP prediction is FATHMM, which could aid in the classification of novel missense SNPs in this gene, providing the identification of mutations potentially associated with drug metabolism.

## Introduction

Intrinsic genetic variations in enzymes and transporters that metabolize drugs can influence the efficacy and toxicity of various drugs (Ahmed, Zhou, Zhou, & Chen, 2016). Different patients may respond differently to the same drug and dose. As a key element in precision medicine, the study of individuals’ responses to medication based on their genomic information allows the evaluation of some specific genetic variants responsible for an individual’s specific response to the drug (Ahmed et al., 2016). Sometimes the effective dose of medication for a particular patient may be fatal or result in therapeutic failure in others, leading to serious adverse effects or even no effect (Ahmed et al., 2016).

In terms of drug metabolism, the monooxygenases from the Cytochrome P450 (CYP) hemeprotein family play a crucial role in metabolizing a wide variety of substances by means of oxidation process (Sridhar, Liu, Foroozesh, & Stevens, 2012). Among such enzymes, the CYP2D6 enzyme represents less than 2% of all CYP enzymes in the liver; nevertheless, this enzyme is involved in the metabolism of endogenous compounds, including hormones and a large number of commonly prescribed exogenous molecules, comprising many antiarrhythmic, β-blockers, neuroleptics, selective serotonin reuptake inhibitors, and tricyclic antidepressants (Brunton, 2006; Sridhar et al., 2012). In addition, the conversion of prodrugs into active compounds or the conversion of drugs into toxic metabolites could involve this enzyme (Brunton, 2006). Thus, CYP2D6 is related to the metabolism of about 25% of currently used drugs, despite the low amount in the liver (Ingelman-Sundberg, 2005b; Ingelman-Sundberg, Sim, Gomez, & Rodriguez-Antona, 2007).

The *CYP2D6* gene is highly polymorphic and the single nucleotide polymorphisms (SNPs) are the most common variations (Ingelman-Sundberg et al., 2007). SNPs are natural gene variations that may have a significant influence on metabolism, clinical efficacy, and side effects in drug therapy, depending on whether the amino acid sequence of the encoded protein is modified and its physiological consequences (Landau, 2005).

Genetic variations are related to dysfunctions in the metabolic capacity of drugs, influencing pharmacokinetics, pharmacodynamics or both, which confer intrinsic differences in drug responses (Ahmed et al., 2016). Therefore, polymorphisms in the CYP2D6 gene could generate four distinct phenotypes based on the metabolizing capacity, including poor, intermediate, extensive and ultra-rapid metabolizers (Arneth, Shams, Hiemke, & Härtter, 2009; Byeon et al., 2018). Hence, CYP2D6 has been considered an important target for pharmacogenomics and pharmacogenetics studies (J. K. Hicks et al., 2013; Ingelman-Sundberg, 2005a; Koski, Ojanperä, Sistonen, Vuori, & Sajantila, 2007).

Currently, the potential impact of missense SNPs has been substantially assessed by means of computational tools. Several approaches have been used for this, because testing these SNPs in the laboratory can be expensive and time-consuming. Therefore, computational tool analysis has become a more accessible approach for preliminary analysis (Shen, Jie D; Deininger Prescott.; Zhao, 2006). Missense SNPs could be classified as deleterious or neutral by a series of *in silico* tools used to predict missense SNP effects on protein function. In general, depending on the strategy used to develop the algorithm, these tools can be classified into four different groups. The first group is the sequence homology-based method, which uses sequence conservation information to assess whether an amino acid substitution affects protein function by means of homologous sequence multiple alignments (Ramensky, 2002). The second group consists of supervised learning methods, where the algorithm “learns” the variants’ conservation patterns and/or physicochemical properties and uses this information to differentiate the SNPs studied by the user as neutral or potentially deleterious (Rodriguez-Casado, 2012). The third group comprises the structure-based methods that evaluate amino acid positions, considering factors such as solvent accessibility and the free energy difference between the wild-type and the mutated amino acids, to analyze the impact of modifications on the protein structure (Gonzalez-Castejon et al., 2011). Finally, the fourth group covers the consensus-based methods, a combination of a variety of methods into a consensus classifier, which could result in significantly improved prediction performance (González-Pérez & López-Bigas, 2011).

Such *in silico* prediction tools have become a target of a number of studies to define the predictive capacity of these tools, in order to indicate the best programs for the evaluation of certain proteins (Arooj et al., 2019; Grimm et al., 2015; Kerr et al., 2017). The predictive capacity of each algorithm is evaluated by statistical measures of performance, which refer to the probability of identifying true deleterious and true neutral mutations (Rodrigues, Santos-Silva, Costa, & Bronze-da-Rocha, 2015). However, the most suitable software to perform the functional prediction of the SNPs present in the *CYP2D6* gene is not yet known. Therefore, in this study we aim to analyze the predictive ability of 20 web-based algorithms of CYP2D6 variants that were phenotypically characterized by *in vivo* and/or *in vitro* studies.

## Material and methods

### Data sets and accession codes

The NCBI Variation Viewer browser (Sayers et al., 2011) was used to access the dbSNP database (build 147). The CYP2D6 Isoform 1 sequence was retrieved with the RefSeq accession code NM_000106.6. The CYP2D6 protein sequence (RefSeq: NP_000097.3) was obtained from NCBI platform and a protein structure was obtained from the Protein Data Bank (PDB ID: 3QM4). Then, to select the validated SNPs, frequency information was verified at the 1000 Genome Project (Altshuler et al., 2012). From this, other databases were used to determine the phenotypes related to drug metabolism associated with the SNPs present in the CYP2D6 enzyme. The Online Mendelian Inheritance in Man (OMIM) database is the primary repository of comprehensive, curated information on genes and genetic phenotypes and the relationships between them (Amberger, Bocchini, Schiettecatte, Scott, & Hamosh, 2015). The Farmacogene Consortium Variation (PharmVar) is a resource for pharmacogenetics and genomics communities, which has information on genes involved in drug metabolism that contribute to drug transport and response (Gaedigk et al., 2018). The SuperCYP database contains information on metabolism and effects on drug degradation of 57 CYPs, allowing the tolerance to drug cocktails to be verified and to find alternative combinations, to efficiently use the metabolic pathways (Preissner et al., 2009). The SuperCYP database contains 1170 drugs with more than 3800 interactions including references (Preissner et al., 2009).

### Frequency data

The *CYP2D6* SNP frequency data were obtained from the 1000 Genomes Project (phase 3) browser (http://www.internationalgenome.org/1000-genomes-browsers). The browser provides the frequencies for all SNPs identified in the genomes, including 2,504 individuals from 26 populations obtained through a combination of low-coverage (2-6x) whole-genome sequence data, targeted deep (50-100x) exome sequencing and dense SNP genotype data. The 26 populations studied were grouped by the predominant ancestry component into five super-populations: African (AFR) (661 individuals), Ad Mixed American (AMR) (347 individuals), East Asian (EAS) (504 individuals), South Asian (489 individuals) and European (EUR) (503 individuals) (Sudmant et al., 2015).

### Conservation analysis

The ConSurf server is a computational tool for estimating the conservation of evolutionary amino acid positions in a protein molecule based on the phylogenetic relations between homologous sequences (Celniker et al., 2013). Using the CYP2D6 3D structure obtained from the Protein Data Bank (PDB ID: 3QM4) (Bernstein et al., 1978; Lauber, Neudecker, Rösch, & Marx, 2003), Consurf carried out a search for close homologous sequences using CSI-BLAST (Angermüller, Biegert, & Söding, 2012) (3 iterations and 0,0001 e-value cut-off) against the UNIREF-90 (Suzek, Huang, McGarvey, Mazumder, & Wu, 2007) protein database. The sequences were then clustered and highly similar sequences removed using CD-HIT (cut-off 95%) (W. Li & Godzik, 2006).

### *In silico* functional analysis of missense SNPs in the *CYP2D6* gene

In order to evaluate the functional effect of the SNPs present in the *CYP2D6* gene, we used 20 computational tools classified into four groups, comprising sequence homology-based methods, supervised learning methods, structure-based methods and consensus-based methods.

#### Sequence homology-based methods

The **Sorting Intolerant From Tolerant** (SIFT) is a multi-step algorithm that uses a sequence homology-based approach to classify amino acid substitutions (Kumar, Henikoff, & Ng, 2009). The analyses were performed under default options.

**PROVEAN** (Protein Variation Effect Analyzer) provides a generalized approach to predict the functional effects of variations on the protein sequence, including single or multiple amino acid substitutions, and insertions and deletions (Choi, Sims, Murphy, Miller, & Chan, 2012). The analyses were performed under default options.

The **Mutation Assessor** evaluates the functional impact of mutations discovered in cancer or nonsense polymorphisms, based on the evolutionary conservation of the amino acids in protein homologs. The analyses were performed as follows: the functional impact score classifies the changes as “low”, “neutral”, which are later considered as neutral, “medium” and “high”, which are classified as deleterious (Reva, Antipin, & Sander, 2011).

The **Panther** tool is composed of two main components: the panther library (panther/lib) and the panther index (panther/x). Panther/lib is a collection of “books,” each representing a protein family as a multiple sequence alignment, a Hidden Markov Model (HMM), and a family tree. Panther/x is an abbreviated ontology for summarizing and navigating molecular functions and biological processes associated with the families and subfamilies (Thomas et al., 2003). The analyses were performed under default options.

**Functional analysis through hidden Markov models** (FATHMM) is a model that uses position-specific information within a multiple sequence alignment (MSA) of homologous sequences, to question the conservation of sequences through the underlying amino acid probabilities modeled by internal correspondence states of several HMMs representing the alignment of homologous sequences and conserved protein domains (Shihab et al., 2013). We used the Inherited Disease option to return predictions capable of discriminating between disease-causing mutations and neutral polymorphisms of Human Phenotype Ontology.

#### Supervised learning methods

**Hansa** uses a set of features (referred to as nsSNP neutral-disease (nsSNPND) discriminatory features) associated with nsSNPs. This nsSNPND discriminatory feature set includes position-specific probabilities, local protein structural status, and the intrinsic properties of the wild-type and mutated residues (Acharya & Nagarajaram, 2012).

**SNAP** (screening for non-acceptable polymorphisms) is a method that combines many sequence analysis tools in a battery of neural networks to predict the functional effects of nsSNPs (Bromberg, Yachdav, & Rost, 2008).

The **SuSPect** (Disease-Susceptibility-based SAV Phenotype Prediction) tool uses the disease-propensity method, which is based on a binomial test comparing the observed numbers of disease-associated and neutral variants in the protein/domain to random expectation. The predictions are pooled following sequences of proteins and with the central chain protein (IBP) (Yates, Filippis, Kelley, & Sternberg, 2014).

**MutPred** is a computational model that relies on SIFT explicitly estimating the probabilities of affecting various structural and functional properties, such as loss of helical propensity, catalytic activity, or post-translational modifications (B. Li et al., 2009).

**PolyPhen-2** (Polymorphism Phenotyping v2) predicts the possible impact of amino acid substitutions on the stability and function of human proteins using functional annotations of missense SNPs, maps that encode SNPs for gene transcripts, extract annotations of protein sequences and structural attributes, and construct profiles of conservation (Adzhubei et al., 2010).

For the five programs, the analyses were performed under default options.

#### Structure-based methods

**Site Directed Mutator** (SDM) is a statistical potential energy function that uses environment-specific amino-acid substitution frequencies within homologous protein families to calculate a stability score, which is analogous to the free energy difference between the wild-type and mutant protein (Pandurangan, Ochoa-Montaño, Ascher, & Blundell, 2017).

**PoPMuSiC** is a tool for the computer-aided design of mutant proteins with controlled thermodynamic stability properties. It evaluates the changes in folding the free energy of a given protein or peptide under point mutations, on the basis of the experimental or modeled protein structure (Dehouck, Kwasigroch, Gilis, & Rooman, 2011).

**HOTMuSiC** is a tool for the computer-aided design of mutant proteins with controlled thermal stability properties. It evaluates the changes in melting temperature of a given protein or peptide under point mutations, on the basis of the experimental or modeled protein structure (Pucci, Bourgeas, & Rooman, 2016).

**Fold-X** is an empirical force field that was developed for the rapid evaluation of the effect of mutations on the stability, folding and dynamics of proteins and nucleic acids (Schymkowitz et al., 2005).

**SNPMusic**, the stability-oriented knowledge-based classifier, uses protein structure, artificial neural networks, and combinations of solvent-dependent statistical potentials to predict whether destabilizing or stabilizing mutations cause disease (Ancien, Pucci, Godfroid, & Rooman, 2018).

All analyses were performed under default options for the five programs.

#### Consensus-based methods

**PredictSNP** is a classifier that uses protein structure, artificial neural networks, and combinations of statistical potentials dependent on solvent accessibility to predict whether destabilizing or stabilizing mutations cause disease (Bendl et al., 2014).

**Condel** is a method to assess the outcome of missense SNVs using a CONsensus DELeteriousness score that combines various tools and integrates their resulting scores by using a WAS-type approach (González-Pérez & López-Bigas, 2011).

**Meta-SNP** is a random forest-based binary classifier to discriminate between disease-related and polymorphic missense SNVs that integrates four existing methods: PANTHER, PhD-SNP, SIFT and SNAP (Capriotti, Altman, & Bromberg, 2013).

The **TransFIC** tool aims to identify somatic point mutations that drive cancer in sequencing projects. The functional impact score classifies the changes as “low”, “medium”, which are later considered neutral, and “high”, which is classified as deleterious (Ben-Hamo & Efroni, 2011; Gonzalez-Perez, Deu-Pons, & Lopez-Bigas, 2012).

**SNPeffect** is a database for phenotyping of single human nucleotide polymorphisms (SNPs) focusing primarily on molecular characterization and annotation of disease variants and polymorphisms in the human proteome (Reumers et al., 2005).

All analyses were performed under default options for the five programs.

### Statistical analysis

The prediction results of the impact of missense *CYP2D6* SNPs using the tools described above were further evaluated using the following statistical measures of performance:

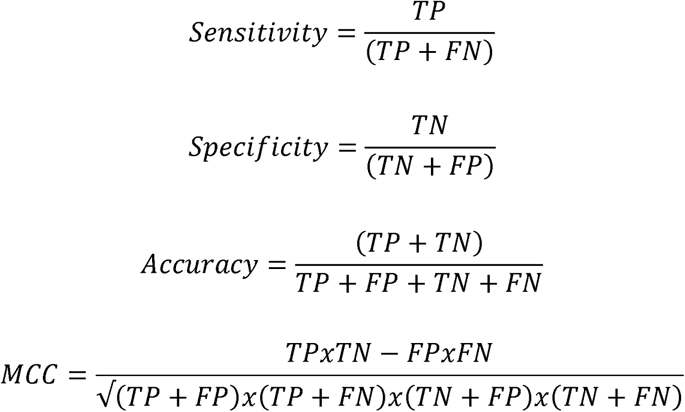

Where TP, TN, FP and FN represent true positives, true negatives, false positives and false negatives, respectively. Sensitivity refers to the probability of identifying true deleterious mutations, whereas specificity represents the probability of identifying true neutral mutations (S. Hicks, Wheeler, Plon, & Kimmel, 2011). The accuracy is the proportion of correct classifications (Berrar, 2018). The MCC is suitable for imbalanced data sets (Boughorbel, Jarray, & El-Anbari, 2017). The Matthews correlation coefficient (MCC) is defined as where −1 indicates perfect negative correlation (the model predicts all negatives as positives, and *vice versa*), 0 indicates no correlation (the model predicts randomly), and +1 indicates perfect positive correlation (the model predicts all real positives as positives and all real negatives as negatives) (Berrar, 2018).

### Combined prediction

In order to improve the performance of the programs, combinations were performed with the best specificity (>70%) and sensitivity (>70%). To determine the result of the combination, two models were adopted: neutral prevails over deleterious (NPOD); and deleterious prevails over neutral (DPON). On the NPOD model, when at least one tool indicated a neutral prediction, the SNP would be considered neutral; and on the DPON model, when at least one tool indicated a deleterious prediction, the SNP would be considered deleterious.

### Receiver operating characteristic

In order to improve the predictive capacity of the FatHMM tool, we used the traditional classification accuracy measures used in medicine, that is, the false positive and true rates (TPR and FPR), also known as sensitivity and specificity, to determine the ideal cutting values (Pepe, Cai, & Longton, 2006). Initially, we determined the minimum (−10.12) and maximum (1.64) scores of the 21 SNPs evaluated. We then evaluated the TPR and FPR using the range of 0.20, gradually increased until reaching the maximum score.

## Results

### Missense SNP selection

From the 328 Missense SNPs available on the NCBI Variation Viewer database (Sayers et al., 2011), a filtering procedure was performed to select only those validated by literature, OMIM, Pharmvar or SuperCYP, selecting those with *in vitro* and/or *in vivo* validation, using this information to identify neutral and deleterious SNPs (Figure 1). After that, 21 missense SNPs were selected (Table 1), of which 6 were neutral and 15 deleterious (Figure 3).

**Fig. 1.**
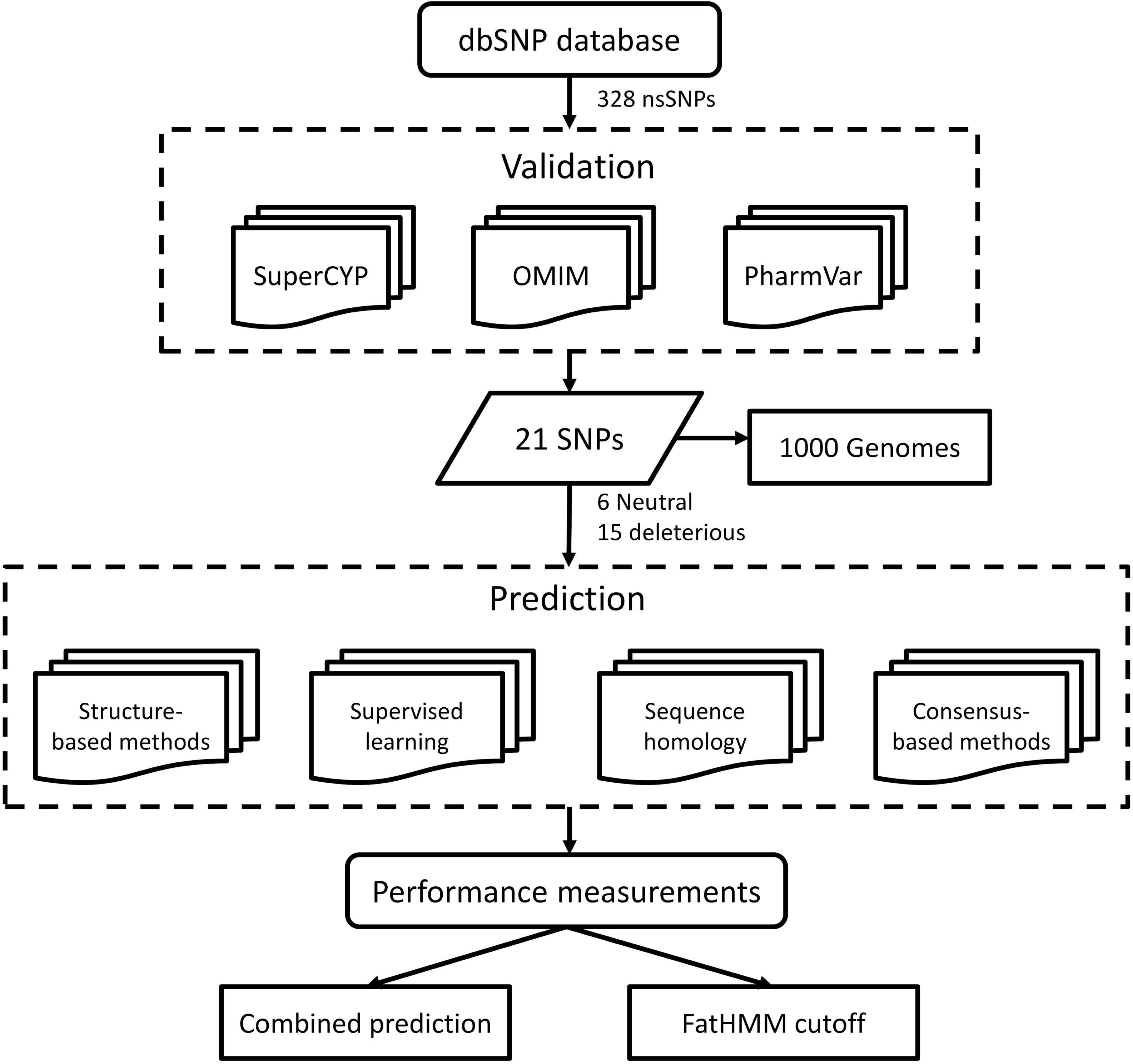
Flowchart used to evaluate the performance measures of nsSNP predictors present in the *CYP2D6* gene. Population frequency information from Project 1000 Genomes was used to select validated SNPs. From this, other databases were used to determine phenotypes related to drug metabolism, associated with SNPs present in the CYP2D6 enzyme. Analysis of the functional effect of the SNPs was carried out using 20 computational tools divided into four distinct groups that were later evaluated by the statistical measures of SEN, SPC, ACC and MCC.

**Table 1.** Information from mutations validated by the literature and the database.

| rsID | Mutation | Metabolism | Literature | Frequency on 1000 Genomes | Population(s) | Reference |
| --- | --- | --- | --- | --- | --- | --- |
| rs769258 | V11M | Poor | Deleterious | 0.017 | AFR, AMR, EAS, EUR, SAS | R. Løvlie et al, 2001 |
| rs1065852 | P34S | Poor | Deleterious | 0.238 | AFR, AMR, EAS, EUR, SAS | Su-Lan Wang et al, 1999 |
| rs373243894 | P41L | Normal | Neutral | 0.001 | AFR | Iris Cohn et al, 2017 |
| rs5030862 | G42R | Poor | Deleterious | ND | ND | Daisuke Tsuzuki et al, 2003 |
| rs118203758 | G42E | Poor | Deleterious | 0.001 | EAS | Daisuke Tsuzuki et al, 2003 |
| rs267608309 | A90V | Normal | Neutral | ND | ND | Sakuyama et al,2008 |
| rs28371706 | T107I | Poor | Deleterious | 0.059 | AFR, AMR, EUR | SuperCYP |
| rs76060075 | G118A | Normal | Neutral | 0.001 | EUR | Chen L. K. et al, 2017 |
| rs569229126 | K147R | Normal | Neutral | 0.001 | EAS | Wang H. et al, 2017 |
| rs267608302 | E156A | Poor | Deleterious | ND | ND | Sakuyama et al,2008 |
| rs5030865 | G169R | Poor | Deleterious | 0.002 | EAS | Su-Lan Wang et al, 1999 |
| rs567606867 | E215K | Increased | Deleterious | 0.001 | AFR, EAS | B Liang et al, 2016 |
| rs371793722 | F219S | Increased | Deleterious | 0.001 | EUR | Ying Su et al, 2016 |
| rs28371717 | A237S | Normal | Neutral | 0.002 | AMR, EAS, EUR, SAS | SuperCYP |
| rs16947 | R296C | Poor | Deleterious | 0.359 | AFR, AMR, EAS, EUR, SAS | (KIM et al, 2013) |
| rs5030867 | H324P | Poor | Deleterious | 0.002 | SAS | B. Evert et al, 1997 |
| rs769157652 | E410K | Normal | Neutral | ND | ND | SuperCYP |
| rs149157808 | E418K | Poor | Deleterious | 0.004 | AFR, AMR, EAS | Joohwan Kim et al, 2013 |
| rs267608319 | R440H | Poor | Deleterious | 0.001 | AMR, EUR | Kathrin Klein, 2007 |
| rs730882171 | R441C | Poor | Deleterious | ND | ND | Kathrin Klein, 2007 |
| rs1135840 | S486T | Increased | Deleterious | 0.401 | AFR, AMR, EAS, EUR, SAS | (KIM et al, 2013) |

ND, not determined; AFR, African; AMR, American; EAS, East Asian; EUR, European; SAS, South Asian;

### SNP frequency in population

CYP2D6 activity varies widely within a population and among ethnically distinct populations (Bernard, 2006). Thus, the 1000 genome was used to determine the frequency of the 21 missense SNPs on the distribution of the five major population groups (Europe, East Asia, South Asia, West Africa and the Americas) (Table 1) (Supp. Figure S1 and S2). The population of East Asia showed the highest prevalence among the SNPs evaluated. Ten SNPs were observed, of which two were neutral (K147R and A237S) and eight deleterious (V11M, P34S, G42E, G169R, E215K, A237S, R296C and E418K). Europe was second with nine SNPs, of which two were neutral (G118A and A237S) and seven deleterious (V11M, P34S, T107I, F219S, R296C, R440H and S486T); then West Africa and America presented eight changes, where seven are deleterious (V11M, P34S, T107I, E215K, R296C, E418K and S486T) and one is neutral in each population (P41L). The lowest number of variations was observed in the South Asian population, where there were only six SNPs, one neutral (A237S) and five deleterious (V11M, P34S, R296C, H324P and S486T). The frequency data for five SNPs (G42R, A90V, E156A, E410K and R441C) could not be determined. Nine SNPs had frequency in more than one population.

### Conservation analysis

The 21 SNPs were mapped by means of ConSurf to identify the degree of conservation of the native residue (Figure 2). This analysis showed that among the missense SNPs, six deleterious ones (T107I, E215K, F219S, R296C, E418K, and S486T) were located in regions considered as variable, three neutral (K147R, A237S and E410K) and five deleterious (V11M, G42R, G42E, G169R and H324P) were located in intermediate regions, and three neutral (P41L, A90V and G118A) and four deleterious (P34S, E156A, R440H and R441C) are located in highly conserved regions with conservation (Figure 2).

**Fig. 2.**
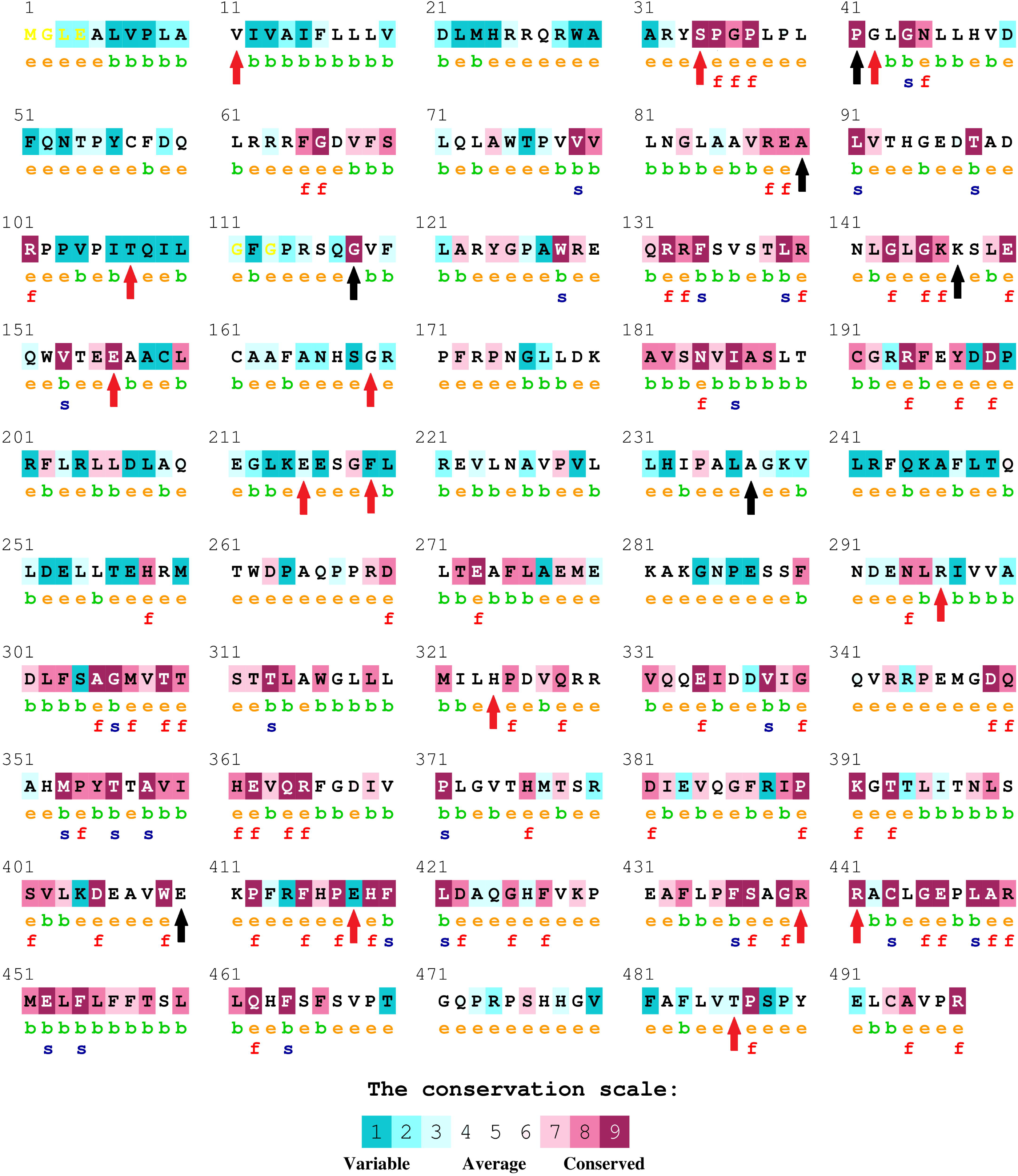
Evolutionary conservation of CYP2D6 amino acid residues obtained from multiple sequence alignment by Consurf. Color intensity increases with degree of conservation. The amino acids are colored based on their conservation grades and conservation levels. A grade of 1 indicates rapidly evolving (variable) sites, which are color-coded in turquoise; 5 indicates sites that are evolving at an average rate, which are colored white; and 9 indicates slowly evolving (evolutionarily conserved) sites, which are color-coded in purple. The deleterious predicted SNPs are marked below the amino acid sequence as red arrows, and the neutral SNPs are marked as black arrows. e - an exposed residue according to the neural-network algorithm; b – a buried residue according to the neural-network algorithm; f – a predicted functional residue (highly conserved and exposed); s - a predicted structural residue (highly conserved and buried).

**Fig. 3.**
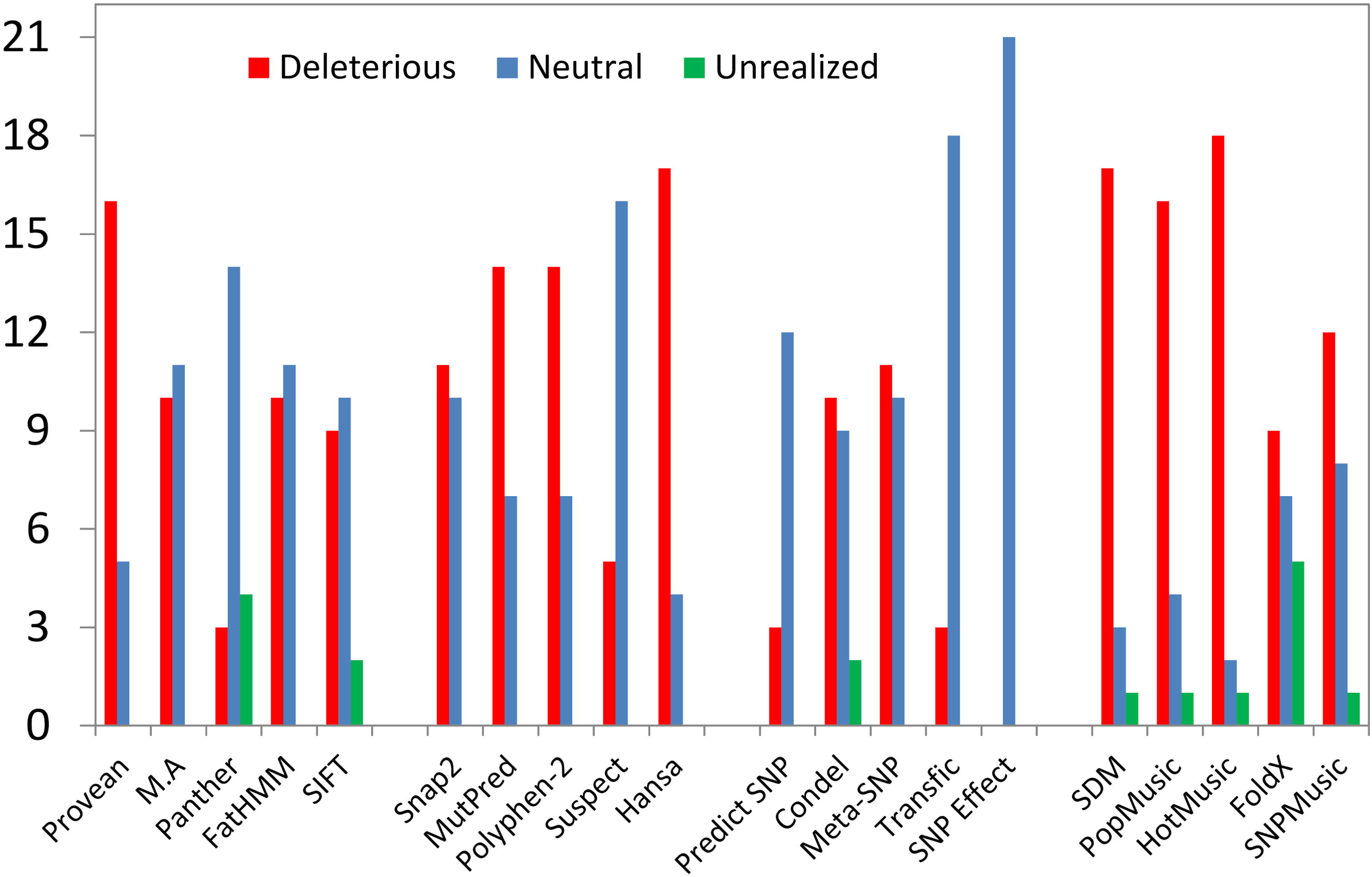
Program performance after impact prediction of the 21 nsSNPs. The red column represents the number of predicted deleterious variations. The green column represents the number of predicted neutral variations. The blue column represents the number of unanalyzed variations.

### Performance measurements

To evaluate the performance of all the tools in this study, we used a set of statistics, such as SEN, SPC, ACC and MCC (Supp. Table S1). For this purpose, the tests were classified as true positive (TP) if the test result corresponded to the deleterious class (pathogenic or harmful) and as true negative (TN) if the result corresponded to the neutral class (neutral or benign). Consequently, a false positive (FP) is a negative test that is classified as positive, and a false negative (FN) is a positive test classified as negative (Grimm et al., 2015). As some data sets are slightly unbalanced, the performance of these predictors was evaluated through the MCC.

Among the groups analyzed, the homology and consensus groups were the ones with the highest number of specific tools, and the learning and structure groups had the highest number of sensitive tools (Figure 4). In general, the consensus group tool set has low sensitivity; structure has low specificity, while homology and machine learning groups present varied results (Figure 4). In the group of methods based on sequence homology, Panther and FatHMM presented specificity greater than 70% and only Provean showed sensitivity >70% (Figure 4). In supervised learning methods, Hansa, Mutpred and Polyphen presented high sensitivity, and only Suspect presented high specificity (Figure 4). The structural methods presented three high sensitivity tools corresponding to SDM, PopMusic and HotMusic (Figure 4). The Consensus-based methods presented one tool with high sensitivity and another with high specificity, which were SNP Effect and Transfic, respectively (Figure 4). PopMusic presented the best accuracy performance compared to other tools (Figure 4). Because these metrics are largely influenced by the size and balance of data sets, MCC was considered the best parameter to measure predictor performance. Considering MCC, which has been described as the best parameter to measure a predictor’s performance (Johnson, Houck, & Chen, 2005), FatHMM achieved the best result among the 20 predictors (0.35).

**Fig. 4.**
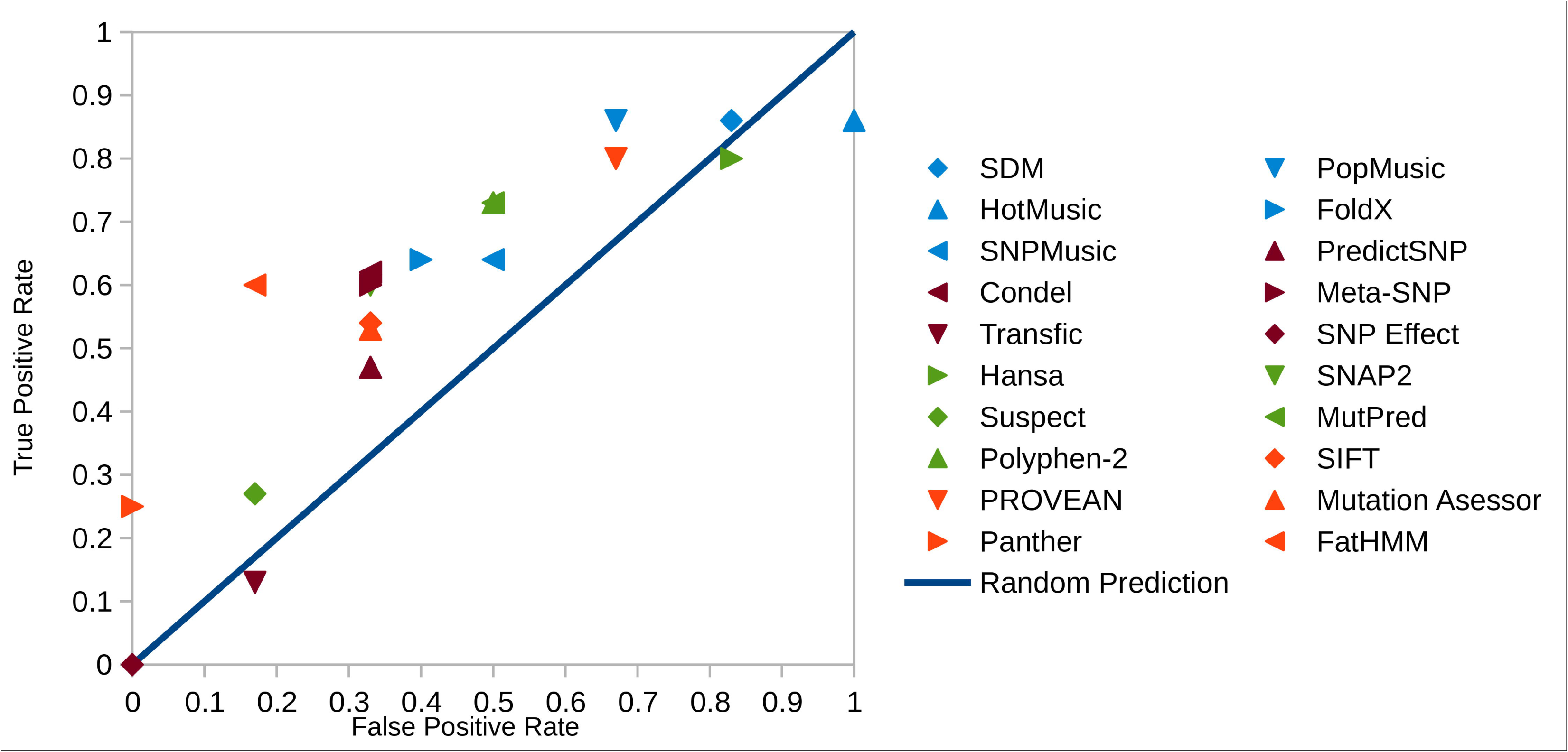
Characteristics of receiver operation of the programs alone. The sequence homology group is represented in orange, the machine learning group is represented in green, the group of consensual methods is represented in brown, and the group based on structure is represented in blue.

### DPON and NPOD Models

The group of low-sensitivity predictors that were combined with the high specificity tools allowed the tools to become more sensitive (Figure 5). There was an increase in the accuracy of the Hansa and SDM servers (Figure 5). In relation to MCC, all the tools that were combined with FatHMM obtained improvement (Figure 5).

**Fig. 5.**
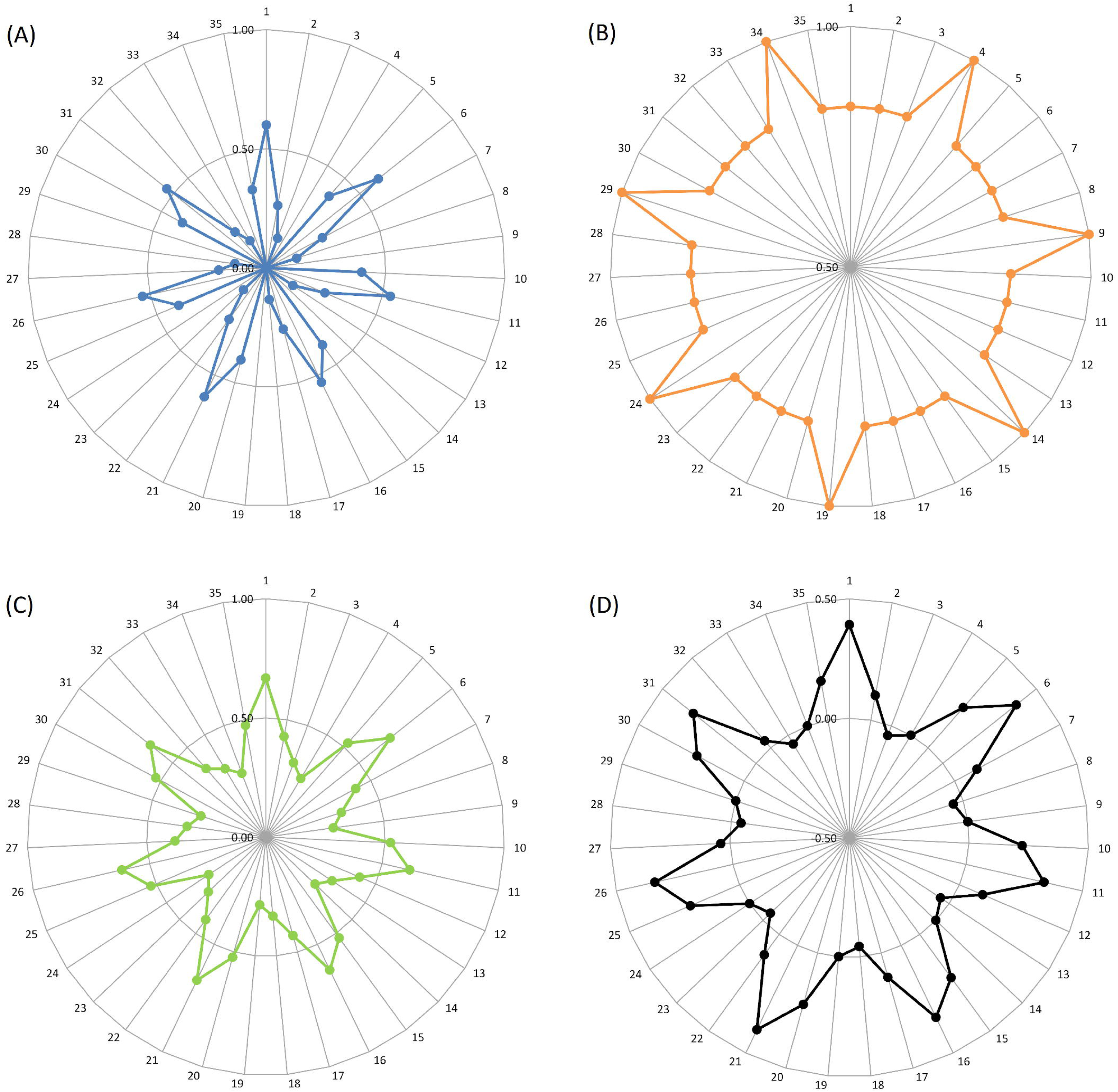
Performance measures of the neutral combination prevail over deleterious (NPOD). The sensitivity is represented in (A), the specificity in (B), the accuracy in (C) and the coefficient of Mathews in (D). The combinations are numbered as follows: (1-5) PROVEAN and (FatHMM; SusPect; Transfic; SNP Effect or Panther, respectively); (6-10) Hansa and (FatHMM; SusPect; Transfic; SNP Effect or Panther, respectively); (11-15) MutPred and (FatHMM; SusPect; Transfic; SNP Effect or Panther, respectively); (16-20) PolyPhen-2 and (FatHMM; SusPect; Transfic; SNP Effect or Panther, respectively); (21-25) SDM and (FatHMM; SusPect; Transfic; SNP Effect or Panther, respectively); (26-30) PopMusic and (FatHMM; SusPect; Transfic; SNP Effect or Panther, respectively); and (31-35) HotMusic and (FatHMM; SusPect; Transfic; SNP Effect or Panther, respectively).

The DPON model obtained the best results, of which all the tools acquired a significant increase in the sensitivity measurements (Figure 6). This improvement was reflected in the determinations of accuracy, which also increased (Figure 6). The combinations between MutPred and Polyphen with FatHMM, Suspect, Transfic or SNPEffect also improved the MCC of this group of predictors (Figure 6).

**Fig. 6.**
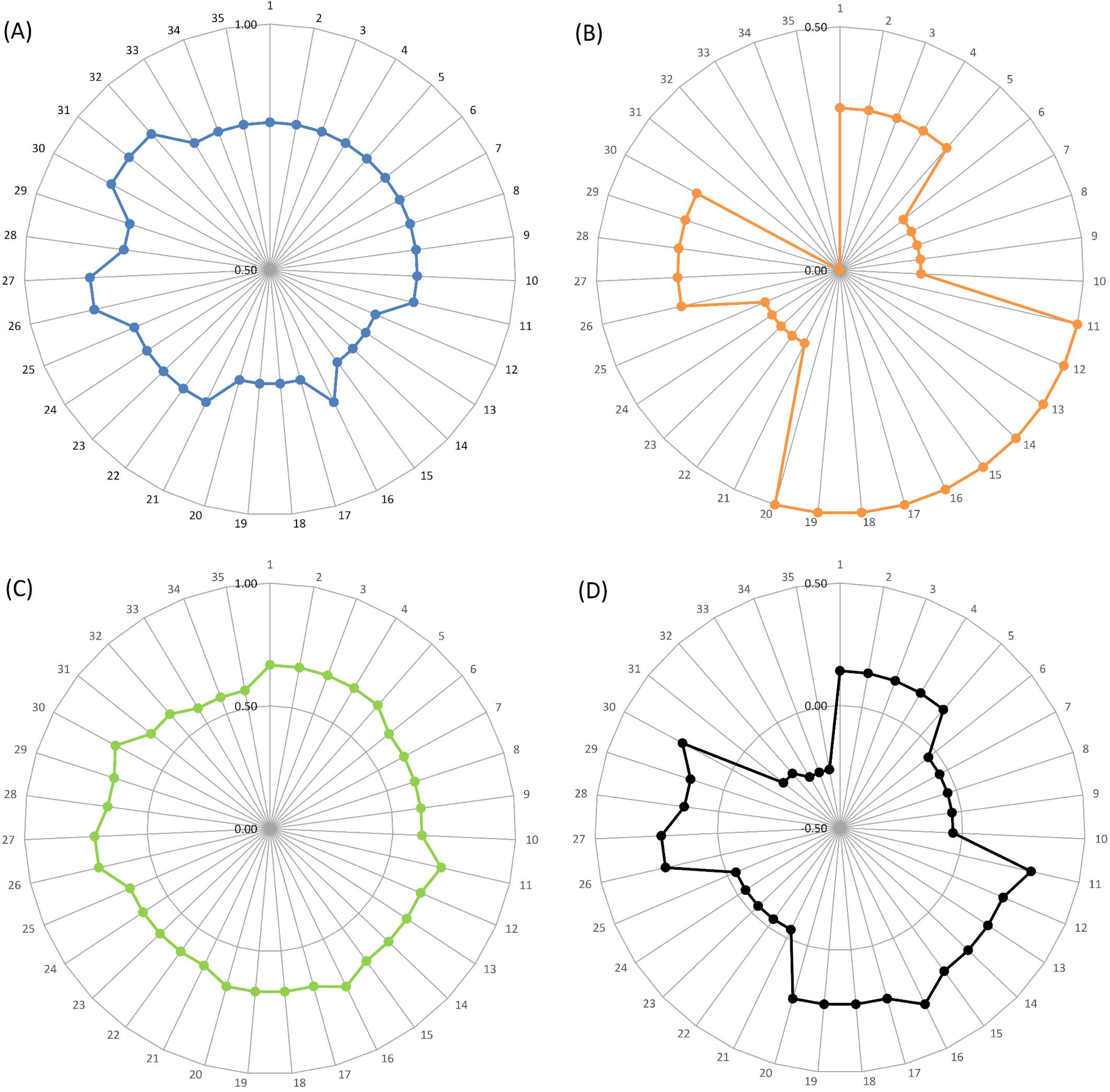
Performance measures of the deleterious combination prevail over neutral (DPON). The sensitivity is represented in (A), the specificity in (B), the accuracy in (C) and the coefficient of Mathews in (D). The combinations are numbered as follows: (1-5) PROVEAN and (FatHMM; SusPect; Transfic; SNP Effect or Panther, respectively); (6-10) Hansa and (FatHMM; SusPect; Transfic; SNP Effect or Panther, respectively); (11-15) MutPred and (FatHMM; SusPect; Transfic; SNP Effect or Panther, respectively); (16-20) PolyPhen-2 and (FatHMM; SusPect; Transfic; SNP Effect or Panther, respectively); (21-25) SDM and (FatHMM; SusPect; Transfic; SNP Effect or Panther, respectively); (26-30) PopMusic and (FatHMM; SusPect; Transfic; SNP Effect or Panther, respectively); and (31-35) HotMusic and (FatHMM; SusPect; Transfic; SNP Effect or Panther, respectively).

### FatHMM Cutoff for CYP2D6

The FatHMM tool demonstrated the best results among the isolated and combined analyses. In order to increase the performance of the tool, we determined a new cutoff, considered −1.15 (Figure 7). Using this strategy, the predictor reached a sensitivity of 80%, accuracy of 76% and MCC of 0.45.

**Fig. 7.**
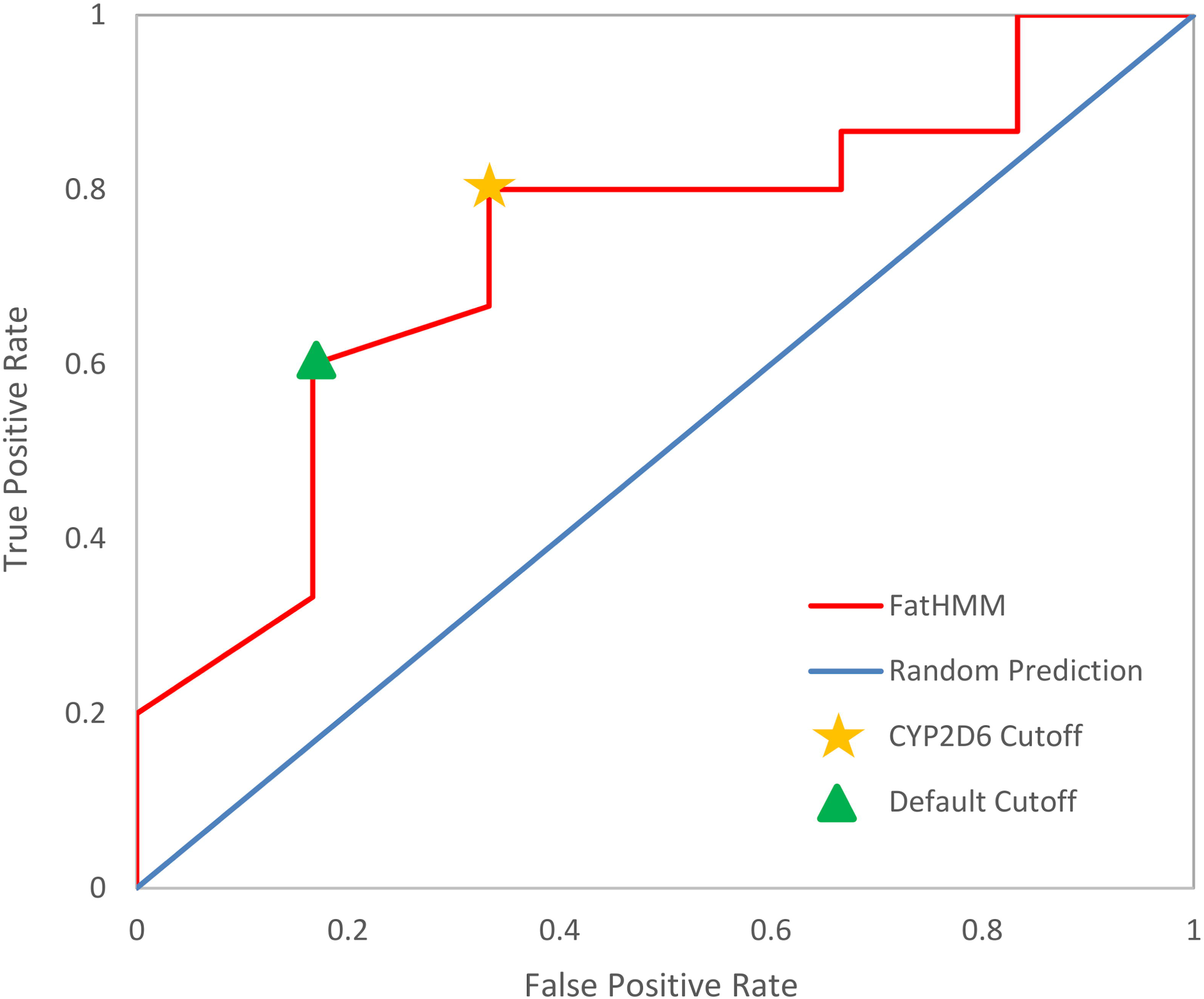
Characteristics of receiver operation of the program FatHMM. Determination of ideal cut values, using the range of 0.02, increasing gradually until reaching the maximum score.

## Discussion

The family of cytochrome P450 enzymes has been the focus of pharmaceutical research for decades, as evidenced by over 100,000 articles in PubMed (Preissner et al., 2009). Despite the vast amount of information on CYP, understanding how the metabolism of this family of enzymes works is still complicated.

The *CYP2D6* gene is highly polymorphic and may include several SNPs at the same time, rather than a single site mutation (Crews et al., 2012). Although the gene is known to have cumulative SNPs, it is possible to determine what a single polymorphism can cause in enzymatic activity. In this study, we selected 21 SNPs previously validated by *in vitro* and/or *in vivo* studies (Table 1), of which 15 single polymorphisms indicated specific alterations where there was an increase, decrease or absence of CYP2D6 enzyme metabolism.

Frequency data validation was obtained from the 1000 Genomes project (Sudmant et al., 2015), where frequency information was obtained from the SNPs of several populations distributed in five groups, of which the East Asian group showed the highest frequency of changes among polymorphisms analyzed (Supp. Figure S2). This information is of great importance for pharmacogenomics to determine responses to drug action in relation to the genetic variations of individuals. To this end, instead of the traditional “one size fits all” approach to drug therapy, the application of targeted drug therapy could allow a more personalized approach, thus minimizing the occurrence of therapeutic failures or adverse effects (Ahasic & Christiani, 2015; Hertz & McLeod, 2016).

The predictive capacity of the set of 20 tools was estimated using statistical measures of SEN, SPC and ACC to evaluate the prediction of the functional significance of the 21 nsSNPs of the *CYP2D6* gene in comparison to the previously reported functional activity. However, the data set presented a number of unbalanced, 15 deleterious and 6 neutral SNPs. Because these metrics are largely influenced by the size and balance of the data sets, the MCC measure has been added because it is best suited for unbalanced data sets. In the evaluated set, the MCC presented results close to zero or negative and only the FatHMM predictor reached an MCC ≥ 0.3, indicating that the systems tested need improvements to predict accurately the impact of missense SNPs on this enzyme. Other independent analyses also reported similar results. In predicting the effect of missense SNPs on *UGT1A1*, Rodrigues et al. (2015) demonstrated that 13 of the 20 tools had an MCC close to zero, and only the predictor of MutPred reached a value of MCC ≥ 0.7 (Rodrigues et al., 2015). Kerr et al. (2017) used 7 tools to predict the functional impact of variants of the *BRCA1*, *BRCA2*, *MLH1* and *MSH2* genes associated with hereditary cancer, and demonstrated that within the evaluated set, 5 algorithms had coefficients below 0.35 and only two tools (Align -GVGD (0.65) and MAPP-MMR (0.59)) had MCCs greater than 0.59 (Kerr et al., 2017).

As described in previous studies, we decided to combine high sensitivity and specificity tools to try to improve the predictive power of these servers (Leong, Stuckey, Lai, Skinner, & Love, 2015). In the combined evaluations, it was possible to see that the combinations between the groups of tools made them more precise, and the best results were observed in the DPON group, due to the data set that presented more deleterious than neutral SNPs. In this last set of analyses, the data generated indicated improvements in the performance of SEN, ACC and MCC. The results suggest that, regardless of the combinations performed, the most appropriate tool for the prediction of SNP impact on CYP2D6 is FatHMM. To improve the predictive capacity of the FatHMM tool, the ROC curve was used to define a new cutoff (−1.15) specific for the *CYP2D6* gene. The concept of an ROC curve is based on the notion of a “separator” (or decision) variable. The frequencies of positive and negative results of the diagnostic test will vary if one changes the “criterion” or “cut-off” for positivity on the decision axis (Hajian-Tilaki, 2013). The new classification threshold increased the measures of SEN, ACC and MCC. Other analyses of independent benchmarks have also reported similar results (Grimm et al., 2015), including a comparative evaluation of 10 tools, distributed in five data sets, to determine which of these tools correctly predicts the pathogenicity of new variants. Among the tools tested, the FatHMM-W demonstrated the best performance in four of the five data sets evaluated (Grimm et al., 2015).

Consurf analysis demonstrated that most SNPs in the *CYP2D6* gene are present in regions considered to be variable or intermediate, which may have contributed to the decrease in accuracy of the tools of the sequence homology group.

The results indicate that the tools have high specificity or high sensitivity, but no precise tools were found. Generally the predictors are constructed to identify changes in several genes. As these tools were not specifically constructed for the *CYP2D6* gene, it is likely that the low accuracy is related to this. Based on this, a tool was built specifically for the CYP, the web server called SCYPPred, which was developed for the prediction of human cytochrome P450 SNPs based on the SVM flanking sequence method ^1^. In order to overcome these limitations, where it is not always possible to obtain specific tools, *in silico* combinational investigations are used to identify deleterious nsSNPs that modify the structure and function of genes (Arooj et al., 2019).

## Conclusion

Although there is a vast range of tools for predicting the functional effect of missense SNPs based on different methodologies, it was not possible to find an accurate tool to analyze these variations in the *CYP2D6* gene. Thus, we proposed the combination of a predictor of high sensitivity and high specificity, seeking to improve the performance of these tools. With the use of more precise tools, it is possible to predict with greater confidence the effect of changes generated in this gene. Although this evaluation showed the weaknesses of the prediction tools currently used, it also demonstrated how to improve them for the development of a more accurate tool, which could aid in the identification of deleterious SNPs potentially associated with deficiencies in the metabolism of the CYP2D6 enzyme present in the human population.

## Acknowledgements

This work was supported by Conselho Nacional de Desenvolvimento Científico e Tecnológico (CNPq), Coordenação de Aperfeiçoamento de Pessoal de Nível Superior (CAPES), Fundação de Apoio a Pesquisa do Distrito Federal (FAPDF) and Fundação de Apoio ao Desenvolvimento do Ensino, Ciência e Tecnologia do Estado de Mato Grosso do Sul (FUNDECT).

## Footnotes

1 The server was not available during the preparation of this manuscript.

## References

Acharya, V., & Nagarajaram, H. A. (2012). Hansa: An automated method for discriminating disease and neutral human nsSNPs. Human Mutation, 33(2), 332–337. https://doi.org/10.1002/humu.21642

Adzhubei, I. A., Schmidt, S., Peshkin, L., Ramensky, V. E., Gerasimova, A., Bork, P., … Sunyaev, S. R. (2010). A method and server for predicting damaging missense mutations. Nature Methods, 7(4), 248–249. https://doi.org/10.1038/nmeth0410-248

Ahasic, A. M., & Christiani, D. C. (2015). Personalized Critical Care Medicine: How Far Away Are We? Seminars in Respiratory and Critical Care Medicine, 36(6), 809–822. https://doi.org/10.1055/s-0035-1564852

Ahmed, S., Zhou, Z., Zhou, J., & Chen, S. Q. (2016). Pharmacogenomics of Drug Metabolizing Enzymes and Transporters: Relevance to Precision Medicine. Genomics, Proteomics and Bioinformatics, 14(5), 298–313. https://doi.org/10.1016/j.gpb.2016.03.008

Altshuler, D. M., Durbin, R. M., Abecasis, G. R., Bentley, D. R., Chakravarti, A., Clark, A. G., … Lacroute, P. (2012). An integrated map of genetic variation from 1,092 human genomes. Nature, 491(7422), 56–65. https://doi.org/10.1038/nature11632

Amberger, J. S., Bocchini, C. A., Schiettecatte, F., Scott, A. F., & Hamosh, A. (2015). OMIM.org: Online Mendelian Inheritance in Man (OMIM®), an Online catalog of human genes and genetic disorders. Nucleic Acids Research, 43(D1), D789–D798. https://doi.org/10.1093/nar/gku1205

Ancien, F., Pucci, F., Godfroid, M., & Rooman, M. (2018). Prediction and interpretation of deleterious coding variants in terms of protein structural stability. Scientific Reports, 8(1). https://doi.org/10.1038/s41598-018-22531-2

Angermüller, C., Biegert, A., & Söding, J. (2012). Discriminative modelling of context-specific amino acid substitution probabilities. Bioinformatics, 28(24), 3240–3247. https://doi.org/10.1093/bioinformatics/bts622

Arneth, B., Shams, M., Hiemke, C., & Härtter, S. (2009). Rapid and reliable genotyping procedure for detection of alleles with mutations, deletion, or/and duplication of the CYP2D6 gene. Clinical Biochemistry, 42(12), 1282–1290. https://doi.org/10.1016/j.clinbiochem.2009.04.009

Arooj, A., Pervez, M. T., Gillani, Z., Chohan, T. A., Tayyab Chaudhry, M., Babar, M. E., & Shah, A. T. (2019). In-Silico Analysis of nsSNPs Associated with CYP11B2 Gene. BioRxiv, 602–615. https://doi.org/10.1101/602615

Ben-Hamo, R., & Efroni, S. (2011). Gene expression and network-based analysis reveals a novel role for hsa-miR-9 and drug control over the p38 network in glioblastoma multiforme progression. Genome Medicine, 3(11). https://doi.org/10.1186/gm293

Bendl, J., Stourac, J., Salanda, O., Pavelka, A., Wieben, E. D., Zendulka, J., … Damborsky, J. (2014). PredictSNP: Robust and Accurate Consensus Classifier for Prediction of Disease-Related Mutations. PLoS Computational Biology, 10(1). https://doi.org/10.1371/journal.pcbi.1003440

Bernard, S. (2006). Interethnic Differences in Genetic Polymorphisms of CYP2D6 in the U.S. Population: Clinical Implications. The Oncologist, 11(2), 126–135. https://doi.org/10.1634/theoncologist.11-2-126

Bernstein, F. C., Koetzle, T. F., Williams, G. J. B., Meyer, E. F., Brice, M. D., Rodgers, J. R., … Tasumi, M. (1978). The protein data bank: A computer-based archival file for macromolecular structures. Archives of Biochemistry and Biophysics, 185(2), 584–591. https://doi.org/10.1016/0003-9861(78)90204-7

Berrar, D. (2018). Performance Measures for Binary Classification. Encyclopedia of Bioinformatics and Computational Biology, 1, 546–560. https://doi.org/10.1016/b978-0-12-809633-8.20351-8

Boughorbel, S., Jarray, F., & El-Anbari, M. (2017). Optimal classifier for imbalanced data using Matthews Correlation Coefficient metric. PLoS ONE, 12(6). https://doi.org/10.1371/journal.pone.0177678

Bromberg, Y., Yachdav, G., & Rost, B. (2008). SNAP predicts effect of mutations on protein function. Bioinformatics, 24(20), 2397–2398. https://doi.org/10.1093/bioinformatics/btn435

Brunton, L. (2006). Summary for Policymakers. Climate Change 2013 - The Physical Science Basis. https://doi.org/10.1017/CBO9781107415324.004

Byeon, J. Y., Kim, Y. H., Lee, C. M., Kim, S. H., Chae, W. K., Jung, E. H., … Lee, Y. J. (2018). CYP2D6 allele frequencies in Korean population, comparison with East Asian, Caucasian and African populations, and the comparison of metabolic activity of CYP2D6 genotypes. Archives of Pharmacal Research, 41(9), 921–930. https://doi.org/10.1007/s12272-018-1075-6

Capriotti, E., Altman, R. B., & Bromberg, Y. (2013). Collective judgment predicts disease-associated single nucleotide variants. BMC Genomics, 14(Suppl 3), S2. https://doi.org/10.1186/1471-2164-14-s3-s2

Celniker, G., Nimrod, G., Ashkenazy, H., Glaser, F., Martz, E., Mayrose, I., … Ben-Tal, N. (2013). ConSurf: Using evolutionary data to raise testable hypotheses about protein function. Israel Journal of Chemistry, 53(3–4), 199–206. https://doi.org/10.1002/ijch.201200096

Choi, Y., Sims, G. E., Murphy, S., Miller, J. R., & Chan, A. P. (2012). Predicting the Functional Effect of Amino Acid Substitutions and Indels. PLoS ONE, 7(10). https://doi.org/10.1371/journal.pone.0046688

Crews, K. R., Gaedigk, A., Dunnenberger, H. M., Klein, T. E., Shen, D. D., Callaghan, J. T., … Skaar, T. C. (2012). Clinical pharmacogenetics implementation consortium (CPIC) guidelines for codeine therapy in the context of cytochrome P450 2D6 (CYP2D6) genotype. Clinical Pharmacology and Therapeutics, 91(2), 321–326. https://doi.org/10.1038/clpt.2011.287

Dehouck, Y., Kwasigroch, J. M., Gilis, D., & Rooman, M. (2011). PoPMuSiC 2.1: A web server for the estimation of protein stability changes upon mutation and sequence optimality. BMC Bioinformatics, 12. https://doi.org/10.1186/1471-2105-12-151

Gaedigk, A., Ingelman-Sundberg, M., Miller, N. A., Leeder, J. S., Whirl-Carrillo, M., & Klein, T. E. (2018). The Pharmacogene Variation (PharmVar) Consortium: Incorporation of the Human Cytochrome P450 (CYP) Allele Nomenclature Database. Clinical Pharmacology and Therapeutics, 103(3), 399–401. https://doi.org/10.1002/cpt.910

Gonzalez-Castejon, M., Marin, F., Soler-Rivas, C., Reglero, G., Visioli, F., & Rodriguez-Casado, A. (2011). Functional Non-Synonymous Polymorphisms Prediction Methods: Current Approaches and Future Developments. Current Medicinal Chemistry, 18(33), 5095–5103. https://doi.org/10.2174/092986711797636081

Gonzalez-Perez, A., Deu-Pons, J., & Lopez-Bigas, N. (2012). Improving the prediction of the functional impact of cancer mutations by baseline tolerance transformation. Genome Medicine, 4(11). https://doi.org/10.1186/gm390

González-Pérez, A., & López-Bigas, N. (2011). Improving the assessment of the outcome of nonsynonymous SNVs with a consensus deleteriousness score, Condel. American Journal of Human Genetics, 88(4), 440–449. https://doi.org/10.1016/j.ajhg.2011.03.004

Grimm, D. G., Azencott, C. A., Aicheler, F., Gieraths, U., Macarthur, D. G., Samocha, K. E., … Borgwardt, K. M. (2015). The evaluation of tools used to predict the impact of missense variants is hindered by two types of circularity. Human Mutation, 36(5), 513–523. https://doi.org/10.1002/humu.22768

Hajian-Tilaki, K. (2013). Receiver Operating Characteristic (ROC) Curve Analysis for Medical Diagnostic Test Evaluation. Caspian Journal of Internal Medicine, 4(2), 627–635. Retrieved from http://www.ncbi.nlm.nih.gov/pubmed/24009950%0Ahttp://www.pubmedcentral.nih.gov/articlerender.fcgi?artid=PMC3755824

Hertz, D. L., & McLeod, H. L. (2016). Integrated patient and tumor genetic testing for individualized cancer therapy. Clinical Pharmacology and Therapeutics, 99(2), 143–146. https://doi.org/10.1002/cpt.294

Hicks, J. K., Swen, J. J., Thorn, C. F., Sangkuhl, K., Kharasch, E. D., Ellingrod, V. L., … Stingl, J. C. (2013). Clinical pharmacogenetics implementation consortium guideline for CYP2D6 and CYP2C19 genotypes and dosing of tricyclic antidepressants. Clinical Pharmacology and Therapeutics, 93(5), 402–408. https://doi.org/10.1038/clpt.2013.2

Hicks, S., Wheeler, D. A., Plon, S. E., & Kimmel, M. (2011). Prediction of missense mutation functionality depends on both the algorithm and sequence alignment employed. Human Mutation, 32(6), 661–668. https://doi.org/10.1002/humu.21490

Ingelman-Sundberg, M. (2005a). Genetic polymorphisms of cytochrome P450 2D6 (CYP2D6): Clinical consequences, evolutionary aspects and functional diversity. Pharmacogenomics Journal, 5(1), 6–13. https://doi.org/10.1038/sj.tpj.6500285

Ingelman-Sundberg, M. (2005b). The human genome project and novel aspects of cytochrome P450 research. Toxicology and Applied Pharmacology, *207*(2 SUPPL.), 52–56. https://doi.org/10.1016/j.taap.2005.01.030

Ingelman-Sundberg, M., Sim, S. C., Gomez, A., & Rodriguez-Antona, C. (2007). Influence of cytochrome P450 polymorphisms on drug therapies: Pharmacogenetic, pharmacoepigenetic and clinical aspects. Pharmacology and Therapeutics, 116(3), 496–526. https://doi.org/10.1016/j.pharmthera.2007.09.004

Johnson, M. M., Houck, J., & Chen, C. (2005). Screening for deleterious nonsynonymous single-nucleotide polymorphisms in genes involved in steroid hormone metabolism and response. Cancer Epidemiology Biomarkers and Prevention, 14(5), 1326–1329. https://doi.org/10.1158/1055-9965.EPI-04-0815

Kerr, I. D., Cox, H. C., Moyes, K., Evans, B., Burdett, B. C., van Kan, A., … Eggington, J. M. (2017). Assessment of in silico protein sequence analysis in the clinical classification of variants in cancer risk genes. Journal of Community Genetics, 8(2), 87–95. https://doi.org/10.1007/s12687-016-0289-x

Koski, A., Ojanperä, I., Sistonen, J., Vuori, E., & Sajantila, A. (2007). A fatal doxepin poisoning associated with a defective CYP2D6 genotype. American Journal of Forensic Medicine and Pathology, 28(3), 259–261. https://doi.org/10.1097/PAF.0b013e3180326701

Kumar, P., Henikoff, S., & Ng, P. C. (2009). Predicting the effects of coding non-synonymous variants on protein function using the SIFT algorithm. Nature Protocols, 4(7), 1073–1081. https://doi.org/10.1038/nprot.2009.86

Landau, R. (2005). Pharmacogenetics: implications for obstetric anesthesia. International Journal of Obstetric Anesthesia, 14(4), 316–323. https://doi.org/10.1016/j.ijoa.2005.03.005

Lauber, T., Neudecker, P., Rösch, P., & Marx, U. C. (2003). Solution structure of human proguanylin: The role of a hormone prosequence. Journal of Biological Chemistry, 278(26), 24118–24124. https://doi.org/10.1074/jbc.M300370200

Leong, I. U. S., Stuckey, A., Lai, D., Skinner, J. R., & Love, D. R. (2015). Assessment of the predictive accuracy of five in silico prediction tools, alone or in combination, and two metaservers to classify long QT syndrome gene mutations. BMC Medical Genetics, 16(1). https://doi.org/10.1186/s12881-015-0176-z

Li, B., Krishnan, V. G., Mort, M. E., Xin, F., Kamati, K. K., Cooper, D. N., … Radivojac, P. (2009). Automated inference of molecular mechanisms of disease from amino acid substitutions. Bioinformatics, 25(21), 2744–2750. https://doi.org/10.1093/bioinformatics/btp528

Li, W., & Godzik, A. (2006). Cd-hit: A fast program for clustering and comparing large sets of protein or nucleotide sequences. Bioinformatics, 22(13), 1658–1659. https://doi.org/10.1093/bioinformatics/btl158

Pandurangan, A. P., Ochoa-Montaño, B., Ascher, D. B., & Blundell, T. L. (2017). SDM: A server for predicting effects of mutations on protein stability. Nucleic Acids Research, 45(W1), W229–W235. https://doi.org/10.1093/nar/gkx439

Pepe, M. S., Cai, T., & Longton, G. (2006). Combining predictors for classification using the area under the receiver operating characteristic curve. Biometrics, 62(1), 221–229. https://doi.org/10.1111/j.1541-0420.2005.00420.x

Preissner, S., Kroll, K., Dunkel, M., Senger, C., Goldsobel, G., Kuzman, D., … Preissner, R. (2009). SuperCYP: A comprehensive database on Cytochrome P450 enzymes including a tool for analysis of CYP-drug interactions. Nucleic Acids Research, 38(SUPPL.1). https://doi.org/10.1093/nar/gkp970

Pucci, F., Bourgeas, R., & Rooman, M. (2016). Predicting protein thermal stability changes upon point mutations using statistical potentials: Introducing HoTMuSiC. Scientific Reports, 6. https://doi.org/10.1038/srep23257

Ramensky, V. (2002). Human non-synonymous SNPs: server and survey. Nucleic Acids Research, 30(17), 3894–3900. https://doi.org/10.1093/nar/gkf493

Reumers, J., Schymkowitz, J., Ferkinghoff-Borg, J., Stricher, F., Serrano, L., & Rousseau, F. (2005). SNPeffect: A database mapping molecular phenotypic effects of human non-synonymous coding SNPs. Nucleic Acids Research, 33(DATABASE ISS.). https://doi.org/10.1093/nar/gki086

Reva, B., Antipin, Y., & Sander, C. (2011). Predicting the functional impact of protein mutations: Application to cancer genomics. Nucleic Acids Research, 39(17). https://doi.org/10.1093/nar/gkr407

Rodrigues, C., Santos-Silva, A., Costa, E., & Bronze-da-Rocha, E. (2015). Performance of In Silico Tools for the Evaluation of UGT1A1 Missense Variants. Human Mutation, 36(12), 1215–1225. https://doi.org/10.1002/humu.22903

Rodriguez-Casado, A. (2012). In silico investigation of functional nsSNPs – an approach to rational drug design. Research and Reports in Medicinal Chemistry, 31. https://doi.org/10.2147/rrmc.s28211

Sayers, E. W., Barrett, T., Benson, D. A., Bolton, E., Bryant, S. H., Canese, K., … Ye, J. (2011). Database resources of the national center for biotechnology information. Nucleic Acids Research, 39(SUPPL. 1). https://doi.org/10.1093/nar/gkq1172

Schymkowitz, J., Borg, J., Stricher, F., Nys, R., Rousseau, F., & Serrano, L. (2005). The FoldX web server: An online force field. Nucleic Acids Research, 33(SUPPL. 2). https://doi.org/10.1093/nar/gki387

Shen, Jie; Deininger Prescott.; Zhao, H. (2006). Applications of computational algorithm tools to identify functional SNPs in cytokine genes. Cytokine, 35(1–2), 62–66.

Shihab, H. A., Gough, J., Cooper, D. N., Stenson, P. D., Barker, G. L. A., Edwards, K. J., … Gaunt, T. R. (2013). Predicting the Functional, Molecular, and Phenotypic Consequences of Amino Acid Substitutions using Hidden Markov Models. Human Mutation, 34(1), 57–65. https://doi.org/10.1002/humu.22225

Sridhar, J., Liu, J., Foroozesh, M., & Stevens, C. L. K. (2012). Insights on cytochrome P450 enzymes and inhibitors obtained through QSAR studies. Molecules, 17(8), 9283–9305. https://doi.org/10.3390/molecules17089283

Sudmant, P. H., Rausch, T., Gardner, E. J., Handsaker, R. E., Abyzov, A., Huddleston, J., … Korbel, J. O. (2015). An integrated map of structural variation in 2,504 human genomes. Nature. https://doi.org/10.1038/nature15394

Suzek, B. E., Huang, H., McGarvey, P., Mazumder, R., & Wu, C. H. (2007). UniRef: Comprehensive and non-redundant UniProt reference clusters. Bioinformatics, 23(10), 1282–1288. https://doi.org/10.1093/bioinformatics/btm098

Thomas, P. D., Campbell, M. J., Kejariwal, A., Mi, H., Karlak, B., Daverman, R., … Narechania, A. (2003). PANTHER: a library of protein families and subfamilies indexed by function. Genome Research, 13(9), 2129–2141. https://doi.org/10.1101/gr.772403

Trynka, G., Hunt, K. A., Bockett, N. A., Romanos, J., Mistry, V., Szperl, A., … Van Heel, D. A. (2011). Dense genotyping identifies and localizes multiple common and rare variant association signals in celiac disease. Nature Genetics, 43(12), 1193–1201. https://doi.org/10.1038/ng.998

Yates, C. M., Filippis, I., Kelley, L. A., & Sternberg, M. J. E. (2014). SuSPect: Enhanced prediction of single amino acid variant (SAV) phenotype using network features. Journal of Molecular Biology, 426(14), 2692–2701. https://doi.org/10.1016/j.jmb.2014.04.026

