## Supplementary material for "Benchmarking analysis of deleterious SNP prediction tools on CYP2D6 enzyme": Supp. Table S1

**Supplementary Tables**

**Supp. Table S1.** Sensitivity (SEN), Specificity (SPC), Matthew Correlation Coefficients (MCC) and Accuracy (ACC) of the twenty predictive tools of SNPs.

| Tools | SEN | SPC | ACC | MCC |
| --- | --- | --- | --- | --- |
| SIFT | 54 | 67 | 58 | 0,19 |
| PROVEAN | 80 | 33 | 67 | 0,14 |
| Mutation Asessor | 53 | 67 | 57 | 0,18 |
| Panther | 25 | 100 | 47 | 0,30 |
| FatHMM | 60 | 83 | 67 | 0,39 |
| Hansa | 80 | 17 | 62 | -0,04 |
| SNAP2 | 60 | 67 | 62 | 0,24 |
| Suspect | 27 | 83 | 43 | 0,11 |
| MutPred | 73 | 50 | 67 | 0,22 |
| Polyphen-2 | 73 | 50 | 67 | 0,22 |
| PredictSNP | 47 | 67 | 52 | 0,12 |
| Condel | 62 | 67 | 63 | 0,26 |
| Meta-SNP | 60 | 67 | 62 | 0,24 |
| Transfic | 13 | 83 | 33 | -0,04 |
| SNP Effect | 0 | 100 | 29 | ND |
| SDM | 86 | 17 | 65 | 0,03 |
| PopMusic | 86 | 33 | 70 | 0,22 |
| HotMusic | 86 | 0 | 60 | -0,22 |
| FoldX | 64 | 60 | 63 | 0,22 |
| SNPMusic | 64 | 50 | 60 | 0,13 |

SEN, sensitivity; SPC, specificity; ACC, accuracy; MCC, Matthews correlation coefficients; ND, not determined;
