## Supplementary material for "Benchmarking analysis of deleterious SNP prediction tools on CYP2D6 enzyme": Supp. Figure S1

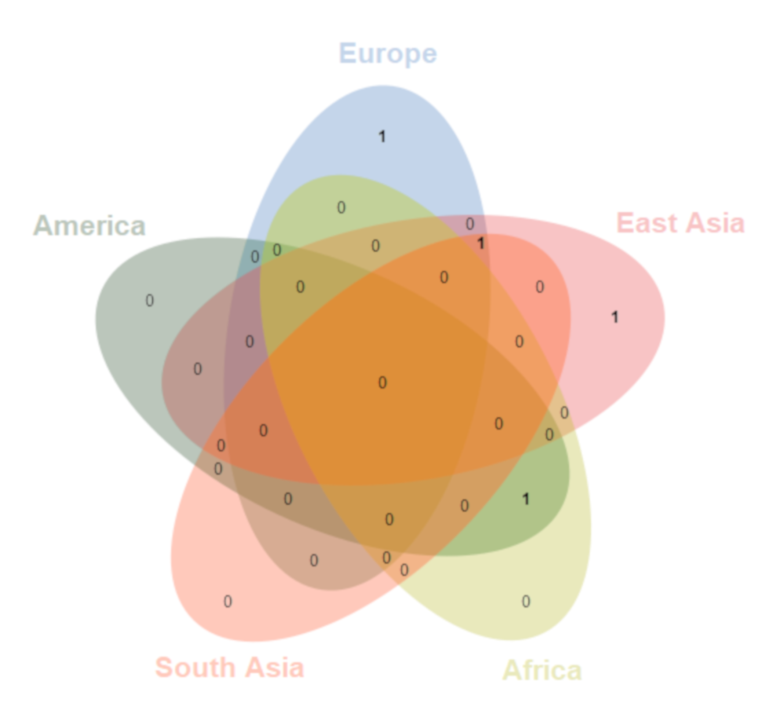

**Supp. Figure S1.** Venn diagram showing the frequency of the neutral SNPs distributed in the five population groups.
