## Supplementary material for "Benchmarking analysis of deleterious SNP prediction tools on CYP2D6 enzyme": Supp. Figure S2

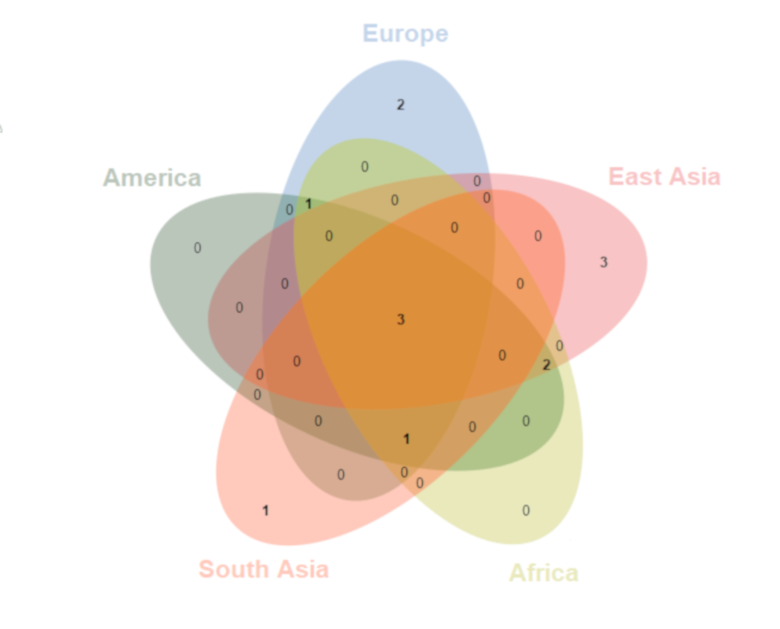

**Supp. Figure S2.** Venn diagram showing the frequency of the deleterious SNPs distributed in the five population groups.
